# An image cryptography method in highly error-prone DNA storage channel

**DOI:** 10.1101/2022.11.08.515529

**Authors:** Xiangzhen Zan, Ranze Xie, Ling Chu, Yanqing Su, Xiangyu Yao, Peng Xu, Wenbin Liu

## Abstract

Rapid development in synthetic technologies has boosted DNA as a potential medium for large-scale data storage. Meanwhile, how to implement data security in DNA storage system is still an unsolved problem. In this paper, we propose an image encryption method based on the modulation-based storage architecture. The key idea is to take advantage of the unpredictable modulation signals to encrypt image in highly error-prone DNA storage channel. Numerical results demonstrate that our image encryption method is feasible and effective with excellent security against various attacks (statistical, differential, noise and data loss, etc.). Compared with other methods by DNA molecules hybridization reaction, the proposed method is more reliable and feasible for large-scale applications.

## 1 Introduction

As the storage medium of genetic information, DNA molecules have the advantage of long durability, high density, and low cost. Recent advancement in synthesis and sequencing technologies has made DNA a promising medium to deal with the challenge of data explosion [1, 2]. Currently, researchers have devoted huge efforts to accurately recover information from the noised sequence pool [3–7]. However, how to ensure the security of private data in DNA storage is an important question still in its infancy.

In 1999, Celland et al. first hid some secret letters in microdots of DNA molecules[8]. Later, Gehani et al. realized the one-time-pad encryption on DNA molecules through DNA microarray technology[9]. In the past decade, researchers continued to explore the encryption potential of complex biochemical processes. In 2014, Yang et al. implemented a 32-bit one-time-pad encryption which simulated one-bit exclusive-or (XOR) operation by DNA strand displacement reaction (SDR) [10]. Later, Peng et al. developed a three-dimensional DNA self-assembly pyramid structure to achieve double-bit encryption[11]. Zhang et al. constructed a DNA origami cryptography method by folding M13 viral scaffolds which could communicate braille-like patterns at nanometer scale [12]. Zakeri et al. accomplished short message communication by chromatogram patterning and multiplexed DNA sequence encoding technology [13]. Peng et al. proposed an one-time-pad cipher algorithm by confusion mapping and random adapter which could guarantee controllable biological security[14]. Recently, some researchers also developed SDR-based chaos system to generate secret keys [15–17]. However, the reliability and practicability of these methods are limited in two aspects. First, the requirement for specially designed and accurately synthesized DNA sequences makes them vulnerable to the complex base errors prevalent in DNA storage. Second, these sophisticated designed experiments may produce unpredictable results in case of some subtle variations in experiment conditions (temperature, time, and ion concentration). And noise environments may even worsen the unpredictability of the results. In addition, these experiment processes are time consuming, difficult to monitor, and not suitable to large-scale applications.

Recently, our group proposed a modulation-based DNA storage architecture which is extremely robust to insertion-deletion-substitution (IDS) errors. The key idea is to transform the binary information to DNA sequences by a modulation signal[18]. Fig. 1 shows an example of the recovered image under different noise levels by three strategies. The first one reconstructs the images directly by multiple sequence alignment (MSA) algorithms. The second one infers a possible modulation signal *M*’ by MSA, and then reconstructs the images using the inferred *M*’ as in Ref. [18]. And the last one recovers images using the true modulation signal *M*. As noises increase, the first two gradually fail to recover the original image while the last one could perfectly recover it. Because the IDS errors are inherent in the synthesis and sequencing processes, the modulation signal could serve as a secure key in a high error DNA storage channel.

**Fig. 1.**
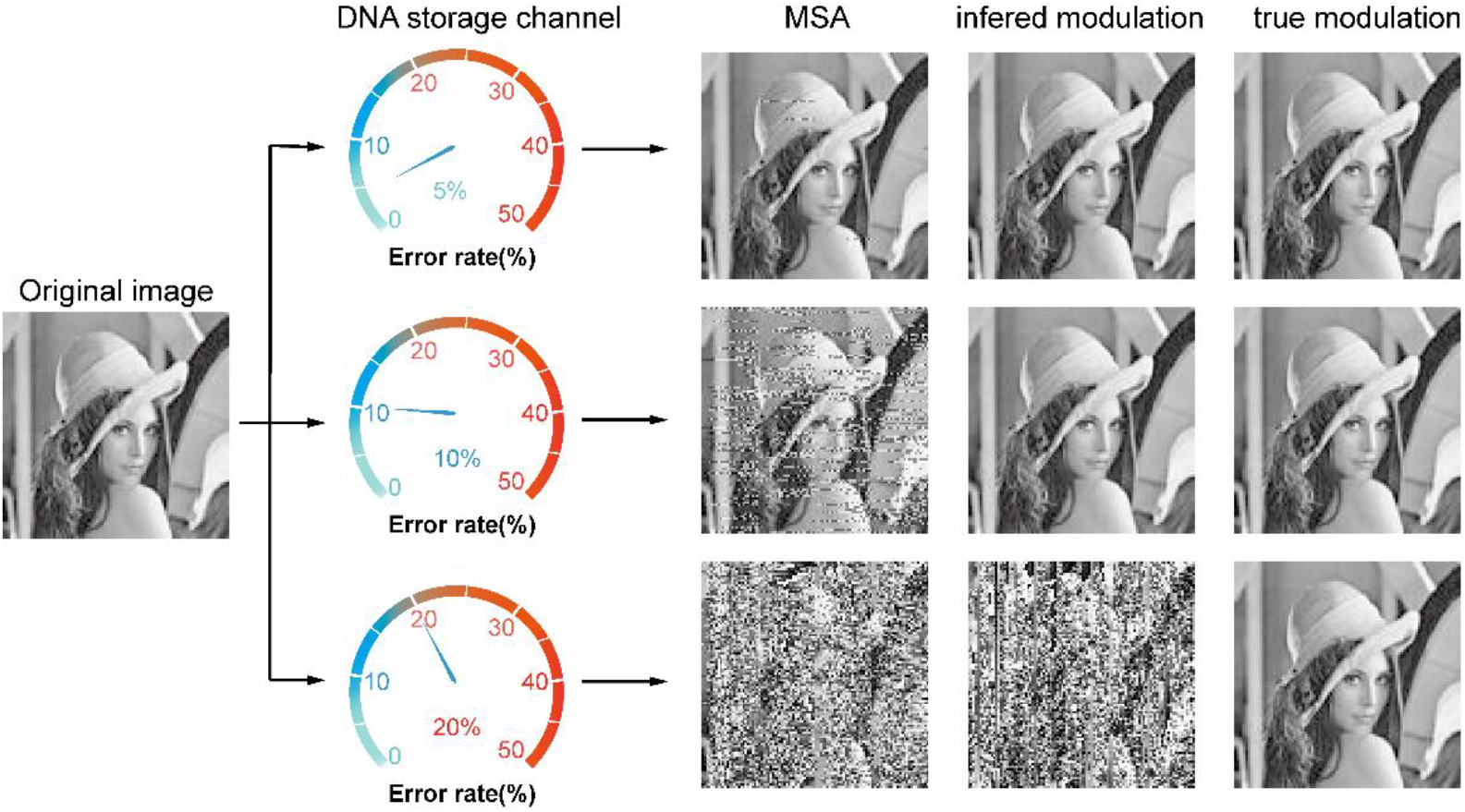
Recovered images at error rate 5%, 10% and 20% by MSA, inferred and true modulation signal. The recovered images by the first two methods gradually become vague as the error rate increases, while the one by the true modulation signal is completely correct.

In this paper, we explore the feasibility of data security in a noisy DNA storage channel. The proposed image encryption scheme consists of two layers: conventional encryption and DNA storage channel encryption. The first layer implements pixel scrambling and diffusion, and the second layer adds further complex confusions in DNA sequences (or DNA pixels) by taking advantage of the uncertainty in DNA storage channel. Simulation results demonstrate that the proposed method could resist cipher attack at DNA sequence level when the noise is larger than 20%. It is also very robust to DNA base errors and sequence losses. Security analysis proves that it has a large key space, is sensitive to the key and plaintext, and can cope with statistical attacks. In sum, the proposed method achieves an excellent combination of the silico-based and carbon-based information security technology and pave a solid foundation for data security in future DNA based information architecture.

## 2 Encryption and decryption

Fig. 2 shows the schematic diagram of the proposed encryption and decryption processes, which include two stages. The first stage performs regular pixel scrambling and diffusion at the binary level. The second stage further encrypts the binary data into DNA sequences by an known modulation key, then they are transmitted through the highly error-prone DNA storage channel, which consists of DNA synthesis, polymerase chain reaction (PCR), and sequencing[3]. Finally, the output ciphertext is a pool of DNA sequences involving large amount of insertion-deletion-substitution errors. The decryption process is the reverse of encryption.

**Fig. 2.**
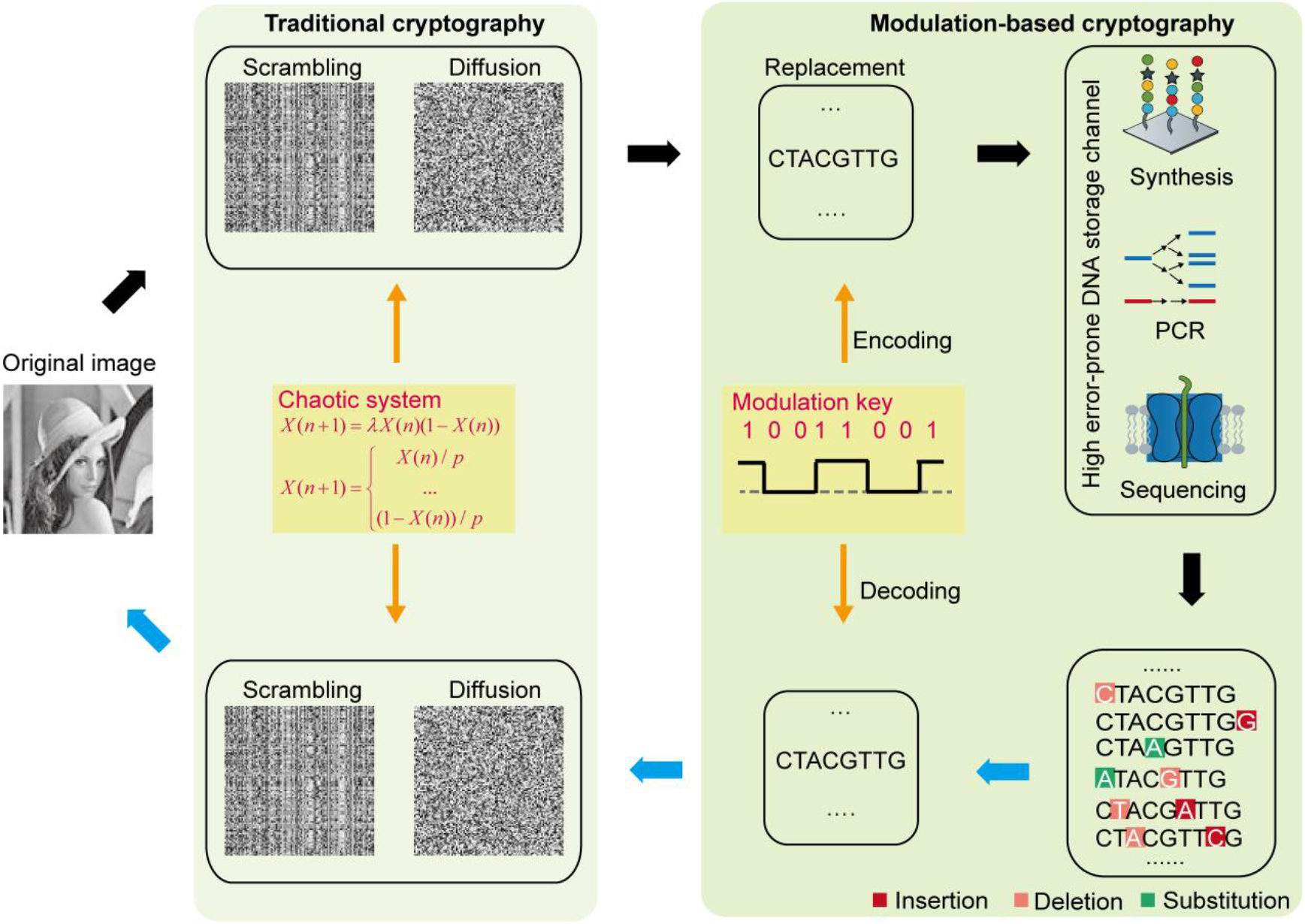
Schematic diagram of the encryption and decryption processes. The left is the traditional cryptography including scrambling and diffusion at the binary level. The right is the modulation-based cryptography at the highly error-prone DNA storage channel. Black arrows represent encryption process while blue arrows indicate decryption process.

### 2.1 Secret key generation

Secret keys mainly consist of two parts. One is the chaotic systems including piecewise linear chaotic map (PWLCM)[19, 20] and logistic map[21, 22]; the other is the modulation sequence.

The dynamic equation of PWLCM can be described by the following function:

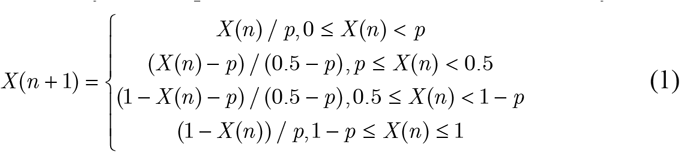

Where the parameter *p* should be in the range of (0, 0.5) and the status value *X*(*n*) is in the range of [0, 1].

The logistic map is defined as follows:

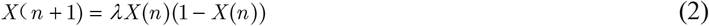

Where the parameter *λ* should be in the range of (0, 4) and the status value *X*(*n*) is in the range of [0, 1].

We use the above chaotic systems to generate three random sequences, two of which are generated by PWLCM with initial status values *X_r_* (0) and *X_c_* (0), and one by logistic map with initial status value *X_d_* (0). For a good encryption process, the initial values are strongly related to the plain image. We use Keccak[23] to hash the plain image to generate a fixed-length value *K* (512 bit), which can be divided into 32 blocks, each of 16-bit. We denote it as *K* = {*k*_1_, *k*_2_,… *k*_32_}. The initial status values are derived as follows:

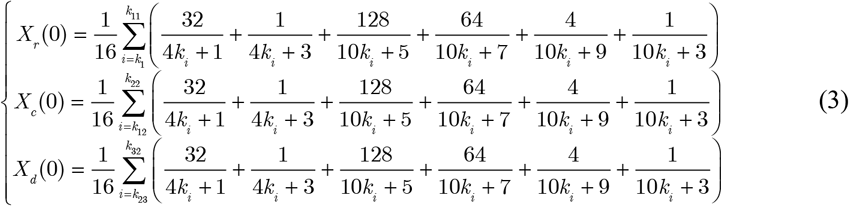

After retrieving the initial value (i.e., *X_r_* (0)) and the corresponding chaotic map, we iterate through the chaotic map *n* times ((i.e., *X_r_*(*n*)) to remove transient processes, and then continue to iterate it to obtain the random sequence of the specified length.

Modulation sequence (key) *M* is a binary sequence where both the percentage of 1s (or 0s) and the consecutive length of 1s (or 0s) can be adjusted as needed, thus ensuring the encoded DNA sequences satisfy biological sequence constraints (i.e., GC balance, no homopolymers). We use *M* to encode the binary data into DNA sequences and correct errors of the sequenced data. The generation of *M* can be referred to Ref. [18].

### 2.2 Encryption algorithm

Suppose the plain image *P* has size of *W* × *H* and the iteration number *n* is in the range of [1000, ∞). Let *N* = *W* × *H*, the detailed encryption process can be depicted as follows.

#### 2.2.1 Traditional cryptography based scrambling and diffusion

**Step 1:** Get the secret keys *λ*, *p*, *X_r_* (0), *X_c_* (0), and *X_d_* (0).

**Step 2:** Use *X_r_*.(0), *X_c_* (0), *n*, and Eq.(1) to generate two sequences of length *W* and *H*, denoted as *S_R_*, *S_C_*. Sort *S_R_* and *S_C_* in ascending order, and perform row-wise and column-wise permutation operations on *P*, respectively, according to the positions of the sorted elements. The scrambled image is denoted as *P*_1_.

**Step 3:** Use *X_d_*(0), *n*, and Eq.(2) to generate a sequence *D* of length of *W*× *H*. Reshape *P*_1_ into one dimensional sequence *Q*. Performing diffusion operation on *Q* using Eq. (4) yields *Q*’. Finally, reshape *Q*’ into a two-dimensional *W* × *H* matrix *P*.

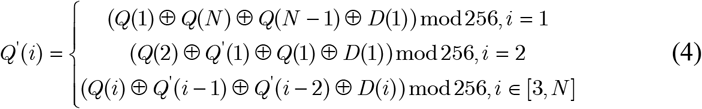

#### 2.2.2 Modulation cryptography based dynamic encryption

**Step1:** Get the secret key *M*.

**Step2:** Transform *P*_2_ into binary form 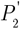, and partition 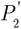 into strands of fixed length *l* (*l* = *len*(*M*)). All these strands are modulated with *M* to generate their corresponding DNA sequences *C* according to a simple mapping rule (00->A, 10->T, 01->C, 11->G). For instance, assuming *M* is ‘**1001**1001**1001**’, the message strand ‘010011010110’ is aligned with *M* into two rows, and a DNA sequence s =‘**CTAC**GTAG**CTTC**’ can be obtained after mapping each column of the two rows into one DNA base.

**Step 3**: Transform *C* into the final cyphertext *C*’ through the error prone DNA storage channel.

### 2.3 Decryption algorithm

The decryption scheme uses the keys including *λ*, *P*, *X_r_* (*n*), *X_c_*(*n*), *X_d_* (*n*), and *M* to execute the reverse operation on the encryption algorithm. First, according to the modulation decoding method[18], *M* is used to correct errors in the sequenced data *C*’ and decode them to obtain the two dimensional pixel matrix *P*_2_. Second, use Eq. (4), Eq. (2), *λ*, and *X_d_* (*n*) to perform reverse diffusion operations on *P*_2_ to get *P*_1_. Finally, use Eq. (1), *p*, *X_r_* (*n*), and *X_c_* (*n*), to do reverse scrambling operations on *p*_1_ to derive the plain image *P*.

## 3 Results

We demonstrate our results on the 100× 100 Lena image as a proof-of-concept. It is encoded by 400 DNA sequences of 200 bases. To investigate the proper noise channel for robust encryption, we take a series of simulation experiments with noises ranging from 2% to 40% and sequence copies ranging from 5 to 10,000.

### 3.1 Key space analysis

The key space of the proposed method is sufficiently large to withstand any brute-force attack. In the traditional decryption process, the receiver needs to know the five parameters *λ*, *p*, *X_r_*(*n*), *X*(*n*) and *X_d_* (*n*). As their valid precision is 10^16^, the key space of the five parameters will be S_Key_ =10^80^ ≈ 2^266^.

Given the sequence length is 200 and the percentage of 1s in the carrier strand is about 0.5, the modulation key space is

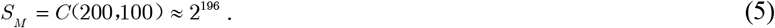

The total key space of our method is

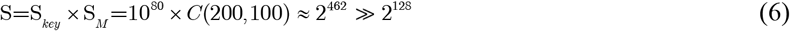

The total key space is much larger than the theoretical secure key value 2^128^ [24]. As the modulation key space alone is larger than 2^128^, we can conclude that the storage channel can serve as another layer for data security.

### 3.2 Cyphertext attack in DNA sequence level

Attackers have two possible ways to decipher the encrypted image in the noisy DNA storage channel. One is to infer a possible modulation key *M*’ by MSA and then decipher the sequenced reads by it, and the other is to directly decipher sequenced reads by the MSA algorithm. Assuming all keys are known except for *M*, we apply one of the famous MSA tools named MAFFT[25] to conduct a series of experiments.

It is impossible to infer a potential modulation key when the error rate is higher than 20%. The attacker may be able to decipher the encrypted image if the inferred key *M*’ is very similar to *M*. Here, we assume that attackers could have sufficient sequence copies to infer *M*. Fig. 3(a) shows the average Hamming distance between *M* and *M*’ increases as the error rate increases. When the error rate is larger than 20%, the average Hamming distance is about 80, and increasing sequence copies may even result in a larger Hamming distance (see the top left corner). Fig. 3(b) further shows the Hamming distance distribution at sequence copies 10,000. The least Hamming distance may reach 32 at error rate 20%. That is, there are at least 32 errors in the inferred modulation keys with 200 bits. As the error rate increases, this lower limit could further increase. Therefore, inferring the true modulation key becomes almost impossible in a high error channel.

**Fig. 3.**
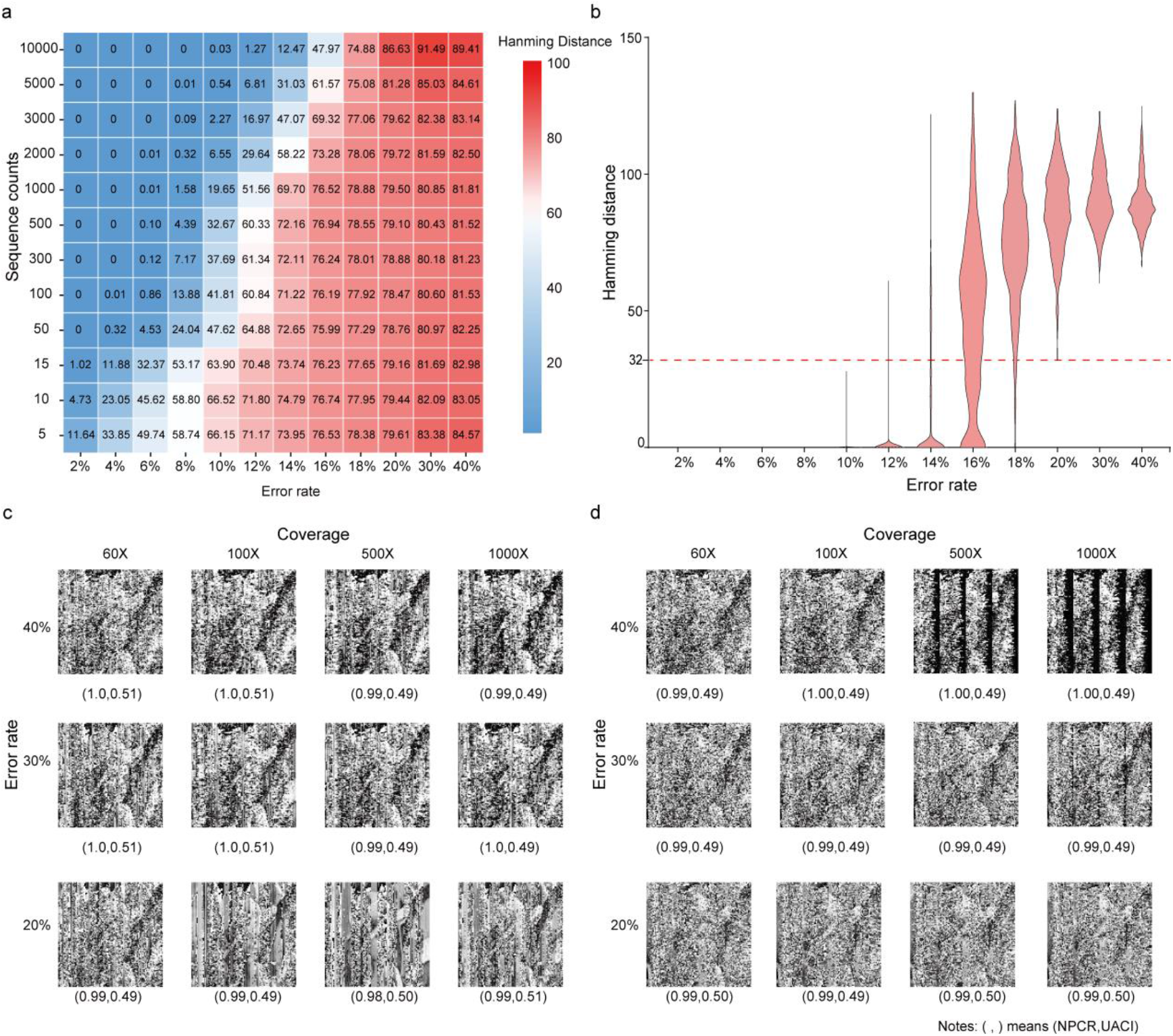
Cipher attack at different error rates and sequence copies. (a) The average Hamming distance between the inferred and true modulation keys. (b) Distribution of the Hamming distances of the inferred modulation keys at sequence copies 10,000. (c) Decrypted images by the inferred modulation keys. (d) Decrypted images by MSA. In (c) and (d), the values in the parentheses denote the NPCR and UACI respectively.

Without knowing the modulation key *M*, it is almost impossible to decipher the real image when the error rate is larger than 20%. To evaluate the difference between the decrypted and the original image, the number of pixels change rate (NPCR) and unified average changing intensity (UACI) are calculated as

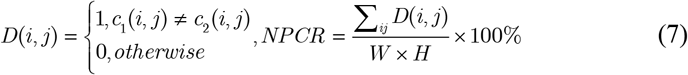

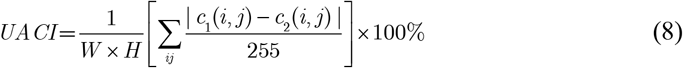

Where *W* and *H* are the width and height of the two images (*c*_1_, *c*_2_). Fig. 3(c) and 3(d) show the decrypted images using different sequence copies by the inferred modulation key and MSA, respectively. Compared with the original image, the decrypted images are all seriously distorted with *UPCR* ≈ 1 and *UACI* ≈ 0.5 even with sequence copies of 1,000. The utilization of traditional cryptographic techniques will further increase crack difficulties.

### 3.3 Sensitivity analysis

The proposed method is sensitive to secrete keys and plaintext. A slight change in the key (i.e., a single bit change) or plaintext could cause a completely different encrypted result. First, the sensitivity of PWLCM map and logistic map has been confirmed in many image-encryption works [19–22]. At the same time, one bit insertion/deletion in the modulation signal will affect the encoding of large amount of pixels. Second, the plaintext sensitivity is accomplished by the pixel diffusion process and the initial status values of the chaotic systems which are strongly related to the plain image.

### 3.4 Statistical analysis

The proposed method can resist statistical attacks. Fig 4. shows the histogram of the pixels in the original image (a) and the encoded 8-base pixel DNA strands (b). Obviously, the distribution of the encoded DNA sequences is flatter than that of the original. Considering the IDS noises in the sequenced reads, the distribution in (b) will tend to be more uniform. Table 1 shows the correlation coefficients of the ciphered image after dislocation and diffusion. All values at the three directions are close to the ideal value of 0[26]. That is, the encrypted pixels are distributed randomly. The information entropy of the cyphered image is 7.950121813, which is very close to the ideal value of 8[26]. Therefore, the encrypted image shows favorable randomness.

**Fig. 4.**
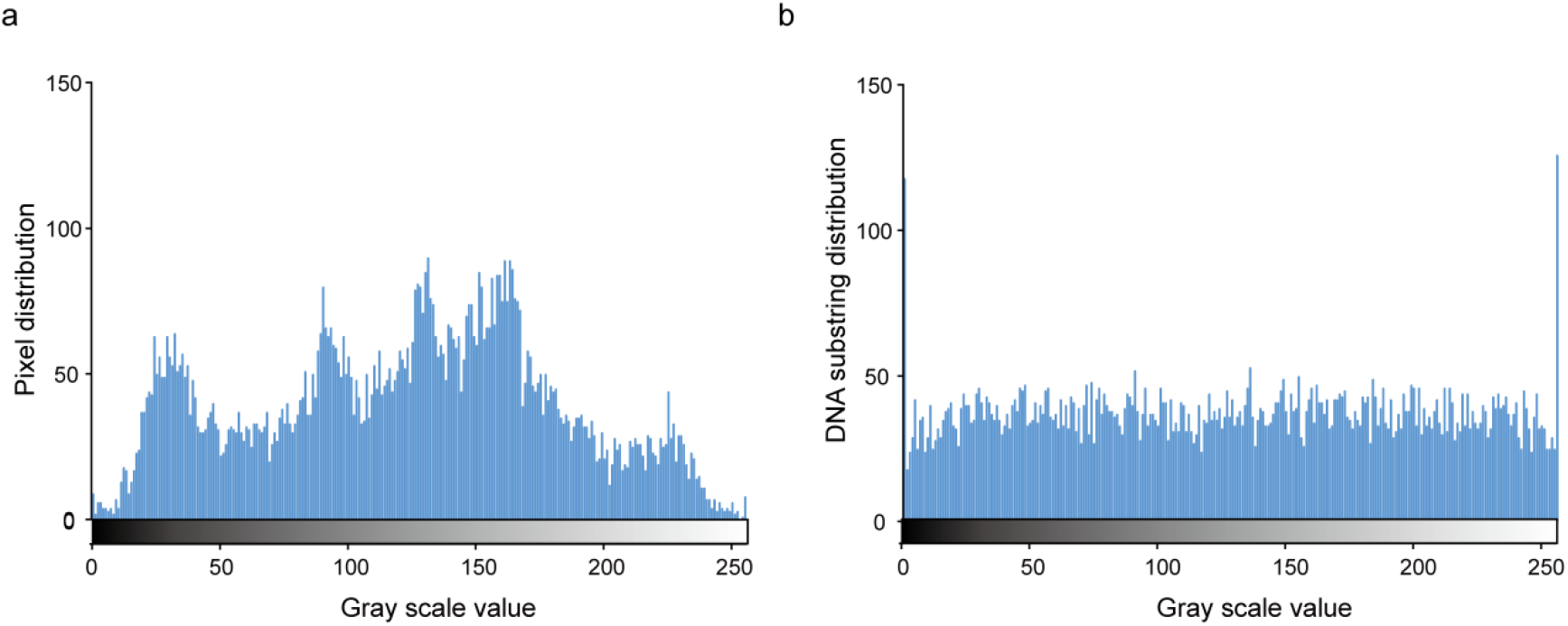
Distribution of pixel intensity histogram. (a) Plain image. (b) Encrypted image.

**Table 1.**
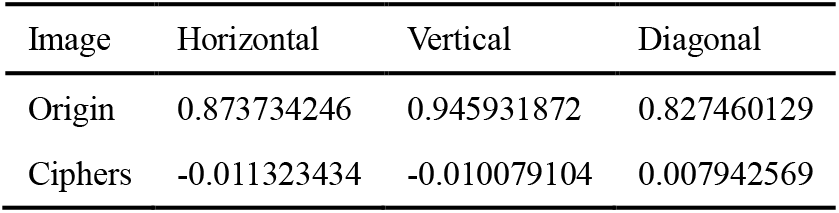
The correlation coefficients in different direction of original and ciphers image

| Image | Horizontal | Vertical | Diagonal |
| --- | --- | --- | --- |
| Origin | 0.873734246 | 0.945931872 | 0.827460129 |
| Ciphers | -0.011323434 | -0.010079104 | 0.007942569 |

### 3.5 Robustness to noises in DNA storage channel

The proposed method is robust to the two most commonly seen errors in DNA storage: base errors and sequence loss. Fig. 5(a) shows the decrypted images at error rate 20~ 40% and sequence copies 50~1000. The original images could be completely deciphered given sufficient sequence copies. Fig. 5(b) shows the decrypted images which could retain the portrait even with loss rate 50%. To the best of our knowledge, such robustness can only be achieved by modulation-based DNA storage architecture [3, 5, 18, 27–29].

**Fig. 5.**
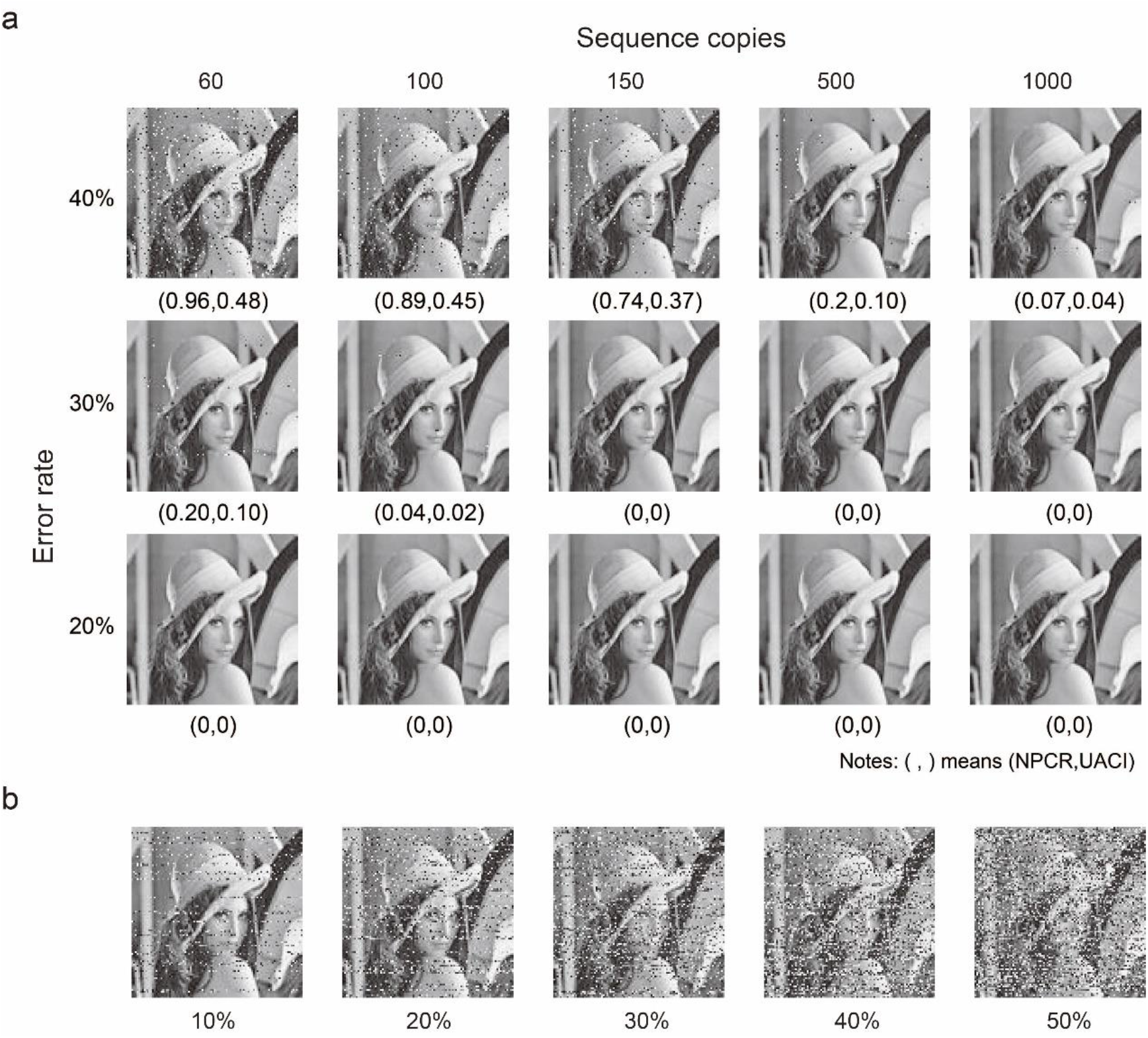
Robustness to base errors and sequence loss. (a) Decryption images at different error rate and sequence copies. (b) Decrypted images at different sequence loss rates

## 4 Comparisons with other encryption methods

Table 2 shows detailed comparisons of existing studies. Compared with other methods, our method has more advantages in terms of encryption using DNA molecules. Firstly, the modulation key and chaotic systems feature our encryption scheme with dynamic encoding and encryption, which can withstand any kinds of brute-force attacks. More importantly, modulation encoding provides a natural way to comply with biochemical constraints for long-term storage. Secondly, its robustness can tolerate extreme environments with high base noises and sequence losses. Thirdly, encrypting data by noise storage channel avoids the complexity and uncertainty in biochemical reactions, such as DNA strand displacement, DNA hiding, and DNA self-assembly. Finally, compared with molecular structure based methods, the proposed method can accomplish higher logical information density (namely 1.0 bits/nt). We believe that all these features endow our method the potential to achieve a reliable, secure, robust, and scalable encryption for DNA storage.

**Table 2.** Comparisons of encryption methods for DNA storage

| Literatures | Dynamic encoding | Dynamic encryption | Robustness | Biological encryption | large scale encryption |
| --- | --- | --- | --- | --- | --- |
| Yang et al[10] | ✓ | * | * | ✓ | × |
| Zakeri et al[13] | × | * | * | ✓ | × |
| Zhang et al[12] | × | * | * | ✓ | × |
| Peng et al[14] | ✓ | ✓ | × | ✓ | × |
| Zhu et al.[17] | ✓ | ✓ | × | ✓ | × |
| This work | ✓ | ✓ | ✓ | ✓ | ✓ |

×- indication of minimum level of support ✓-Indication of acceptable level of support \*-Partial fulfillment

## 5 Conclusions

We propose an image encryption method for DNA storage, which can be divided into two parts: conventional encryption and DNA storage channel encryption. The proposed method highlights the importance of the unpredicted modulation signals in a highly error-prone DNA storage channel. Simulation results show that our method is feasible and effective for encrypting and decrypting images when the error rate of the DNA storage channel is higher than 20%. Further analysis of the security shows that it is sensitive to both keys and plaintexts, has a large enough key space, and can resist various attacks (i.e., statistical, only cipher text, noise and data loss, etc.). Compared with other state-of-the-art encryption methods, our approach has high logical information density, compliance with biochemical constraints, and strong robustness to base errors and sequence loss, and is thus more suitable for large-scale DNA encryption storage. Though designed for image encryption, our method can also be applicable to other areas of encryption. Relying on the powerful error correction capability of the modulation-based DNA storage architecture, we believe our approach will further accelerate the arrival of large-scale DNA encrypted storage.

## Acknowledgements

This work was supported by the National Natural Science Foundation of China (grant nos.62072128 and 62002079).

## Notes

### Competing Interest Statement

The authors have declared no competing interest.

## References

[1] L. Qian, Q. Ouyang, Z. Ping, F. Sun, and Y. Dong, “DNA storage: research landscape and future prospects,” National Science Review, vol. 7, no. 6, pp. 1092–1107, 2020.

[2] L. C. Meiser, B. H. Nguyen, Y.-J. Chen, J. Nivala, K. Strauss, L. Ceze, and R. N. Grass, “Synthetic DNA applications in information technology,” Nature Communications, vol. 13, no. 1, pp. 352, 2022/01/17, 2022.

[3] P. L. Antkowiak, J. Lietard, M. Z. Darestani, M. M. Somoza, W. J. Stark, R. Heckel, and R. N. Grass, “Low cost DNA data storage using photolithographic synthesis and advanced information reconstruction and error correction,” Nature Communications, vol. 11, no. 1, pp. 5345, 2020/10/22, 2020.

[4] L. C. Meiser, P. L. Antkowiak, J. Koch, W. D. Chen, A. X. Kohll, W. J. Stark, R. Heckel, and R. N. Grass, “Reading and writing digital data in DNA,” Nat Protoc, vol. 15, no. 1, pp. 86–101, Jan, 2019.

[5] W. H. Press, J. A. Hawkins, S. K. Jones, J. M. Schaub, and I. J. Finkelstein, “HEDGES error-correcting code for DNA storage corrects indels and allows sequence constraints,” Proceedings of the National Academy of Sciences of the United States of America, vol. 117, no. 31, pp. 18489–18496, Aug 4, 2020.

[6] J. Jeong, S. J. Park, J. W. Kim, J. S. No, H. H. Jeon, J. W. Lee, A. No, S. Kim, and H. Park, “Cooperative Sequence Clustering and Decoding for DNA Storage System with Fountain Codes,” Bioinformatics, Apr 27, 2021.

[7] Y. Erlich, and D. Zielinski, “DNA Fountain enables a robust and efficient storage architecture,” Science, vol. 355, no. 6328, pp. págs. 950–954, 2017.

[8] C. T. Clelland, V. Risca, and C. Bancroft, “Hiding messages in DNA microdots,” Nature, vol. 402, pp. 750–750, 12/16, 1999.

[9] A. Gehani, T. LaBean, and J. Reif, “DNA-based Cryptography,” Aspects of Molecular Computing: Essays Dedicated to Tom Head, on the Occasion of His 70th Birthday, N. Jonoska, G. Păun and G. Rozenberg, eds., pp. 167–188, Berlin, Heidelberg: Springer Berlin Heidelberg, 2004.

[10] J. Yang, J. Ma, S. Liu, and C. Zhang, “A molecular cryptography model based on structures of DNA self-assembly,” Chinese Science Bulletin, vol. 59, no. 11, pp. 1192–1198, 2014/04/01, 2014.

[11] W. Peng, D. Cheng, and C. Song, “One-time-pad cryptography scheme based on a three-dimensional DNA self-assembly pyramid structure,” PLOS ONE, vol. 13, no. 11, pp. e0206612, 2018.

[12] Y. Zhang, F. Wang, J. Chao, M. Xie, H. Liu, M. Pan, E. Kopperger, X. Liu, Q. Li, J. Shi, L. Wang, J. Hu, L. Wang, F. C. Simmel, and C. Fan, “DNA origami cryptography for secure communication,” Nature Communications, vol. 10, no. 1, pp. 5469, 2019/11/29, 2019.

[13] B. Zakeri, P. A. Carr, and T. K. Lu, “Multiplexed Sequence Encoding: A Framework for DNA Communication,” PLoS One, vol. 11, no. 4, pp. e0152774, 2016.

[14] W. Peng, S. Cui, and C. Song, “One-time-pad cipher algorithm based on confusion mapping and DNA storage technology,” PLoS ONE, vol. 16, no. 1, pp. e0245506, 2021.

[15] C. Zou, X. Wei, Q. Zhang, C. Zhou, and S. Zhou, “Encryption Algorithm Based on DNA Strand Displacement and DNA Sequence Operation,” IEEE Transactions on NanoBioscience, vol. 20, no. 2, pp. 223–234, 2021.

[16] Y. Wang, Z. Li, and J. Sun, “Three-Variable Chaotic Oscillatory System Based on DNA Strand Displacement and Its Coupling Combination Synchronization,” IEEE Transactions on NanoBioscience, vol. 19, no. 3, pp. 434–445, 2020.

[17] E. Zhu, X. Luo, C. Liu, and C. Chen, “An Operational DNA Strand Displacement Encryption Approach,” Nanomaterials, 12, 2022].

[18] X. Zan, R. Xie, X. Yao, P. Xu, and W. Liu, “A robust and efficient DNA storage architecture based on modulation encoding and decoding,” bioRxiv, pp. 2022.05.25.490755, 2022.

[19] M. Alawida, A. Samsudin, J. S. Teh, and R. S. Alkhawaldeh, “A new hybrid digital chaotic system with applications in image encryption,” Signal Processing, vol. 160, pp. 45–58, 2019/07/01/, 2019.

[20] P. Zhou, J. Du, K. Zhou, and S. Wei, “2D mixed pseudo-random coupling PS map lattice and its application in S-box generation,” Nonlinear Dynamics, vol. 103, no. 1, pp. 1151–1166, 2021/01/01, 2021.

[21] L. Sui, B. Liu, Q. Wang, Y. Li, and J. Liang, “Color image encryption by using Yang-Gu mixture amplitude-phase retrieval algorithm in gyrator transform domain and two-dimensional Sine logistic modulation map,” Optics and Lasers in Engineering, vol. 75, pp. 17–26, 2015/12/01/, 2015.

[22] L. Sui, K. Duan, J. Liang, Z. Zhang, and H. Meng, “Asymmetric multiple-image encryption based on coupled logistic maps in fractional Fourier transform domain,” Optics and Lasers in Engineering, vol. 62, pp. 139–152, 2014/11/01/, 2014.

[23] G. Bertoni, J. Daemen, M. Peeters, and G. Van Assche, “Keccak,” in ADVANCES IN CRYPTOLOGY - EUROCRYPT 2013, 2013, pp. 313–314.

[24] Y. Dong, G. Zhao, Y. Ma, Z. Pan, and R. Wu, “A novel image encryption scheme based on pseudo-random coupled map lattices with hybrid elementary cellular automata,” Information Sciences, vol. 593, pp. 121–154, 2022/05/01/, 2022.

[25] K. Katoh, K. Misawa, K. i. Kuma, and T. Miyata, “MAFFT: a novel method for rapid multiple sequence alignment based on fast Fourier transform,” Nucleic Acids Research, vol. 30, no. 14, pp. 3059–3066, 2002.

[26] H. M. Ghadirli, A. Nodehi, and R. Enayatifar, “An overview of encryption algorithms in color images,” Signal Processing, vol. 164, pp. 163–185, 2019/11/01/, 2019.

[27] S. R. Srinivasavaradhan, S. Gopi, H. Pfister, and S. Yekhanin, “Trellis BMA: coded trace reconstruction on IDS channels for DNA storage,” 2021.

[28] S. M. H. T. Yazdi, R. Gabrys, and O. Milenkovic, “Portable and Error-Free DNA-Based Data Storage,” Scientific Reports, vol. 7, Jul 10, 2017.

[29] L. Song, F. Geng, Z.-Y. Gong, X. Chen, J. Tang, C. Gong, L. Zhou, R. Xia, M.-Z. Han, J.-Y. Xu, B.-Z. Li, and Y.-J. Yuan, “Robust data storage in DNA by de Bruijn graph-based de novo strand assembly,” Nature Communications, vol. 13, no. 1, pp. 5361, 2022/09/12, 2022.

